# *Drosophila*–virus genotype interactions dominate transmission, virulence and load, eroding additive fitness variance

**DOI:** 10.64898/2026.05.21.726842

**Authors:** Aleksei Belyi, Yiyang Du, Alastair J Wilson, Ben Longdon, Francis M Jiggins

## Abstract

Host–parasite coevolution is expected to generate strong selection for susceptibility and infectivity that has the potential to erode genetic variation. Despite this, natural populations often retain extensive genetic variation in these traits. Negative frequency-dependent selection could explain the maintenance of variation since it results in genotype-by-genotype interactions on fitness that, unlike additive genetic variance among hosts and pathogens, is masked from selection and so can remain ‘cryptic’. Here we combine coevolutionary modelling with large-scale experimental infection assays to quantify how host–pathogen interactions structure fitness variance in a vertically transmitted virus system, *Drosophila melanogaster* and sigma virus. Simulations show that coevolution typically erodes additive genetic variance in host and pathogen fitness, concentrating variance in host–virus interaction terms. Consistent with these predictions, experiments spanning 90 host–virus genotype combinations reveal that transmission, viral load and virulence are overwhelmingly governed by host–virus genetic interactions rather than host or virus main (i.e., additive) effects. As a result, neither host resistance alleles nor viral genotypes confer consistently higher fitness across genetic backgrounds. Interactions tend to be sex-specific, further limiting heritable fitness variation. Our results demonstrate that coevolution can substantially mask heritable genetic variation from selection by rendering fitness context dependent. This extensive cryptic genetic variation may be revealed either when ecological or evolutionary conditions shift, or when the process of coevolution itself alters the direction of selection. This demonstrates the need to account for genotype-specific interactions when forecasting evolutionary responses.

## Introduction

Hosts are harmed by their parasites and pathogens (hereafter parasites), while parasites must infect their host to survive [1]. This antagonistic conflict can drive rapid evolution, with change in one partner leading to opposing adaptations in its adversary. This coevolutionary battle can cause immune systems to change and diverge [2], favour sexual over asexual reproduction [3] and drive sexual selection [4].

This rapid evolution is fuelled by high levels of genetic variation in host susceptibility to infection, which have been observed across organisms from bacteria to humans [5–11]. Similarly, parasites exhibit genetic variation in infectivity [12,13]. Although the exact mechanisms maintaining this variation remain the matter of debate [1], in both hosts and parasites coevolution is thought to play a key role [14,15]. Alleles conferring resistance may be costly because they have harmful pleiotropic effects on other traits, or because alleles that confer resistance to one pathogen genotype confer susceptibility to other genotypes. When alleles that increase resistance spread through a population, the prevalence of the pathogens targeted by that allele can decline to a point where these costs outweigh the benefits of resistance [16]. The equivalent process can occur in the parasite population, resulting in negative frequency-depended selection (NFDS). Importantly, this form of selection can maintain genetic variation in both parties [17].

The importance of coevolution as a driver of evolutionary change will depend on the extent to which this genetic variation in host susceptibility and pathogen infectivity results in genetic variation in fitness [13]. It is common for infectious disease to be a major determinant of host survival and reproduction, and the ability of pathogens to infect and transmit is strongly affected by host genotype [12,18–20]. However, this may not translate into effects on (average) allelic fitness if there are non-additive interactions between host and pathogen genotypes. For instance, a host allele that has high fitness when exposed to one pathogen genotype may not be beneficial overall if it has low fitness when exposed to a different genotype or when uninfected. Similarly, genetic lineages of pathogens may pass through different host genotypes where they have high and low fitness. The premise that genotype-by-genotype interactions on fitness can enable NFDS and so maintain global genetic variance is recognised by game theoretic and quantitative genetic treatments of coevolutionary dynamics across many different types of biological interaction [21,22]. NFDS can lead to stable equilibrium allele frequencies—at which point all host and pathogen alleles have the same expected fitness [23]. This means there is no additive genetic variation for fitness among hosts or pathogens at equilibrium. Nevertheless, the outcome of each infection still depends on non-additive genetic factors–specifically the interaction of specific host and pathogen genotypes. However, time lags in evolutionary responses mean that coevolutionary models of host–parasite systems often do not predict stable equilibria, but instead generate temporal fluctuations in the direction of selection and therefore allele frequencies [16]. By preventing the population from reaching a stable equilibrium, these temporal cycles in selection can potentially move genetic variance back into main effects, restoring genetic variance in fitness.

The sigma virus (DMelSV) of *Drosophila melanogaster* is a natural pathogen that infects 0–20% of wild flies [24–26]. DMelSV is transmitted vertically through eggs and at a lower rate through sperm [24,27–31], and laboratory and field studies have estimated that infection results in a ∼20% reduction in fitness [24,32,33]. In *D. melanogaster* populations there is considerable genetic variation in viral titres and transmission [24]. Resistant alleles of three genes—*p62* (*ref(2)P*) [5,34], *CHKov1* [35], and *Ge-1* [36]—reduce both viral replication and transmission, together explaining 59% of the genetic variance in viral load [15]. The effects of these alleles on transmission differ between males and females [5,37]. There is also genetic variation in the viral population, with some viral genotypes escaping the effects of the *p62* resistance allele [38–40]. The spread of the *p62* resistance allele has driven the evolution of DMelSV viral types capable of infecting resistant flies [26,39]. This viral type appears to be recent [40]; European DMelSV populations containing both types shared a common ancestor only ∼200 years ago [25]. Its rapid spread was observed in France and Germany during the 1980s [41], where it largely replaced the viral type permissive/sensitive to *p62* over about a decade.

The observation that DMelSV carries a substantial cost to the host is at first sight surprising as vertically transmitted parasites rely on host survival and reproduction for their own transmission. A classical explanation is that there is a trade-off between transmission rate (proportion offspring infected) and virulence (number of offspring), mediated by viral load,[42–50]. Therefore, increasing viral replication may enhance transmission fidelity while simultaneously reducing host fitness. Consequently, viral replication provides a mechanism by which the evolutionary interests of host and parasite become decoupled, generating an intrinsic conflict that can promote reciprocal coevolutionary dynamics [51–57].

To investigate the role of host-genotype–by–virus-genotype interactions, we combined modelling with empirical experiments of *D. melanogaster* and DMelSV. We find that host and viral genotypes strongly influence transmission, viral load and virulence. However, consistent with coevolutionary predictions [17], much of this variance reflects pronounced genotype–by–genotype interactions. Thus, despite extensive genetic effects on infection outcomes, coevolution has reduced heritable variation in fitness for both host and pathogen, potentially limiting evolutionary responses.

## Results

### Coevolution creates genetic interactions that reduce fitness variance

To examine coevolutionary dynamics in a vertically transmitted virus, we simulated a host population segregating a resistance allele and a viral population segregating for an escape allele that is insensitive to/can overcome host resistance, as observed for DMelSV in nature. In line with classical gene-for-gene models, both host resistance and viral escape incurred costs [58]. We used parameter values within the empirical ranges estimated for natural *Drosophila*–sigma virus populations [24,29–33], with resistance cost (for which no field estimate is available) varied across a broad plausible range. The system most commonly produced oscillatory coevolutionary dynamics, of two types: damped oscillations converging on a stable polymorphism in host and virus (Fig 1A–1C, S2A–S2C, S3A–S3C), or persistent cycles in allele frequencies in both populations (Fig 1D–1F, S2D–S2F, S3D–S3F). If there were no costs of resistance and escape, there was a partial selective sweep of host resistance which stalled when the viral escape allele was fixed (S4A Fig).

**Fig 1.**
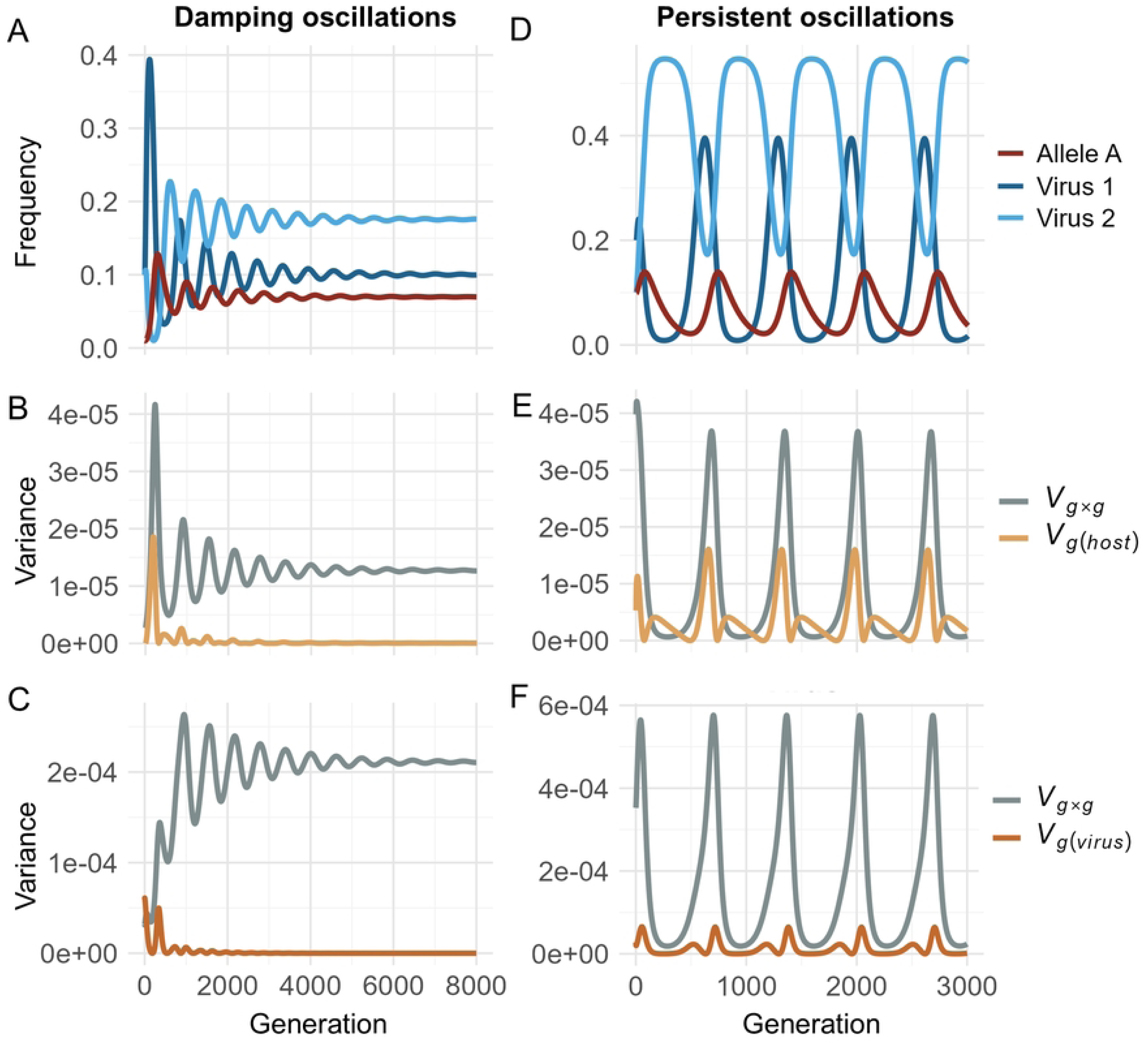
Time series of coevolutionary dynamics between a host and vertically transmitted pathogen under a gene-for-gene model. The host carries one biallelic resistance locus (A/a), where allele A is dominant and partially resistant to the virus. The virus has two genotypes, where Virus 1 is susceptible to host allele A and Virus 2 is an escape variant that is not affected. The left column (A–C) illustrates damped oscillations converging on a stable polymorphism (*t₁* = 0.68, *t₂* = 0.66, *c* = 0.23, *r* = 0.27, *d* = 0.006). The right column (D–F) shows persistent allele-frequency cycles (*t₁* = 0.72, *t₂* = 0.70, *c* = 0.255, *r* = 0.27, *d* = 0.007). Parameters are *t*, vertical-transmission rate; *c*, viability cost of infection; *r*, proportional reduction in virus transmission in resistant hosts; *d*, per-locus fertility cost of carrying a dominant resistance allele. (B, E) Host-fitness genetic variance partitioned into the additive host component *V_G(host)_* (orange) and the host × virus interaction *V_G×G_* (grey). (C, F) Viral-fitness variance partitioned analogously into *V_G(virus)_* (red) and *V_G×G_* (grey). The data supporting these figures can be found in S9 Data. To investigate these dynamics more systematically, we explored a range of parameter values (Fig 2; full parameter sweeps in S6 and S7 Fig). The coefficient of variation of viral-genotype frequencies (Fig 2A) tracks the dynamical regime, with a lower-amplitude corresponding to damped oscillations approaching a stable equilibrium. In these regions of parameter space, the fitness variance was concentrated in the interaction term, with the proportion of additive genetic variance approaching zero for both host and virus (*V_G_* / (*V_G_* + *V_G×G_*; Fig 2B and C). Conversely, a high coefficient of variation reflects persistent or unstable cycles, which maintained additive genetic variance in fitness (Fig 2 B and C). However, even under these conditions the interaction variance (*V_G×G_*) typically remained the largest single component. Thus, genotype-by-genotype interactions arising during coevolution are expected, in most conditions, to substantially depress additive genetic variance in host and virus fitness.

To link these dynamics to fitness, we decomposed the genetic variance in fitness into additive components for host and virus, *V_G(host)_* and *V_G(virus)_*, and a genotype-by-genotype term, *V_G×G_*, capturing genetic effects on infection outcomes that do not translate into heritable fitness (for the host, this term also includes differences in the fitness of infected and uninfected individuals). With damped oscillations (Fig 1A), additive genetic variances *V_G(host)_* and *V_G(virus)_* decline towards zero as equilibrium was approached, while *V_G×G_* remains high (Fig 1B and 1C). This is an expected signature of NFDS eroding any net advantage of individual genotypes. In contrast, stable cycles (Fig 1D) maintained additive genetic variance in fitness—*V_G(host)_* and *V_G(virus)_*—though time (Fig 1E and 1F). However, *V_G×G_* typically dominated for both host and viral fitness, although this oscillated through time along with the genotype frequencies. These qualitative patterns were robust across different genetic interactions, such as matching-allele and multi-locus gene-for-gene models (S2 and S3 Fig; middle and bottom). When there are selective sweeps of viral escape alleles and partial sweeps of host resistance, *V_G×G_* becomes important for the host once viral escape becomes common (S4B and S4C Fig).

In other regions of parameter space, polymorphisms were not maintained so coevolution ceased (loss of one or both virus genotypes or loss of host resistance). Oscillations occurred predominantly at basal transmission rates of roughly 67 to 83% and infection costs of about 1% to 30%, ranges that overlap empirical estimates of transmission and virulence in wild *Drosophila*–sigma virus populations (S6 and S7 Fig; detailed parameter ranges, outcome classifications, and a broader exploration of parameter space are provided in S1 Text) [24,33,59].

### Experimental data confirms genetic variation in sigma virus fitness is strongly reduced by G_H_×G_P_ interactions

The fitness of a vertically transmitted pathogen is determined by the number of offspring produced by infected hosts and the rate at which the virus is transmitted to those offspring. We quantified how host and viral genotypes influence sigma virus transmission rates (Fig 3A and 3B) by infecting nine *D. melanogaster* lines from the *Drosophila* Genetic Reference Panel (DGRP) [60,61] with ten strains of sigma virus. As the virus is vertically transmitted, we crossed viruses into the fly lines (via backcrossing with balancers), ensuring natural transmission dynamics [31]. This crossing design generated all 90 combinations independently, decoupling viral isolate from host genetic background.

**Fig 2.**
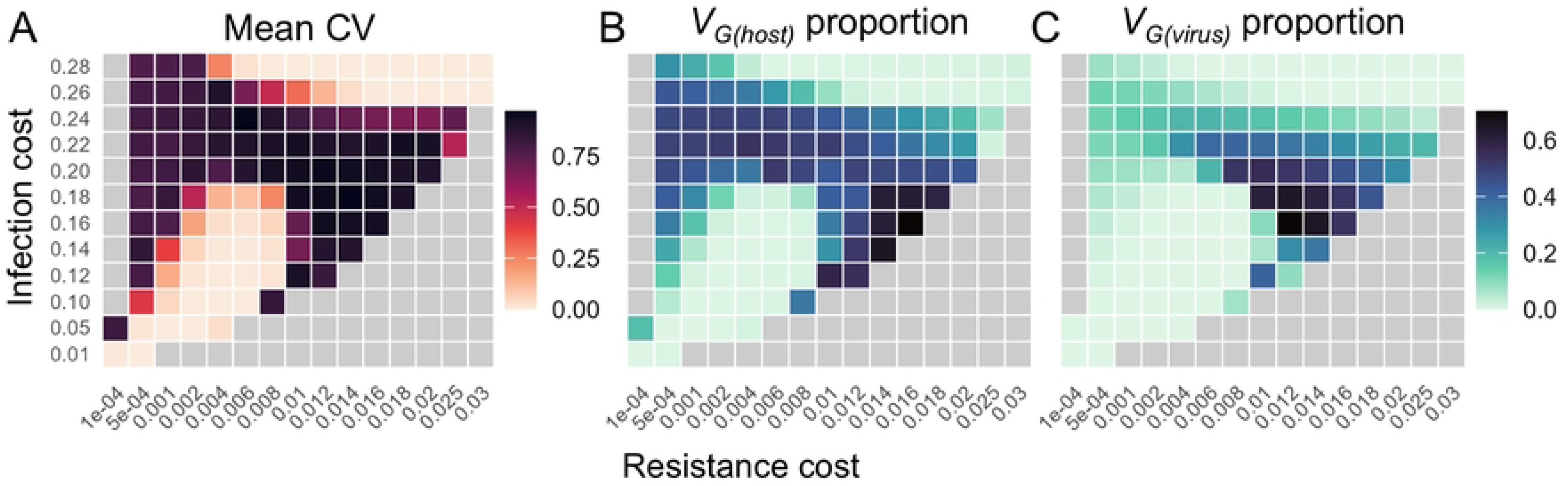
Coevolutionary dynamics and fitness variance across parameter space. Coevolution was simulated assuming a gene-for-gene model over a range of per-locus fertility costs of resistance *d* and viability cost of infection *c*. Baseline transmission rates were fixed at *t₁* = 0.73, *t₂* = 0.70 (within the empirically observed sigma virus range); other parameters are the same as in Fig 1. Each cell is a 40,000-generation run with statistics computed over the final 20,000 generations. (A) The mean coefficient of variation (CV) of viral-genotype frequencies, averaged across the two viral strains. (B) The proportion of additive genetic variance in host fitness, *V*_G(host)_ / [*V*_G(host)_ + *V*_G×G_]; and (C) the corresponding proportion for viral fitness. Variance components in (B) and (C) were restricted to generations with viral prevalence > 0.01%. Parameter combinations leading to the loss of either both viruses or just allele A are shown as grey cells. Full parameter space with outcome classification is given in S6 Fig and S7 Fig. The data supporting these figures can be found in S9 Data.

**Fig 3.**
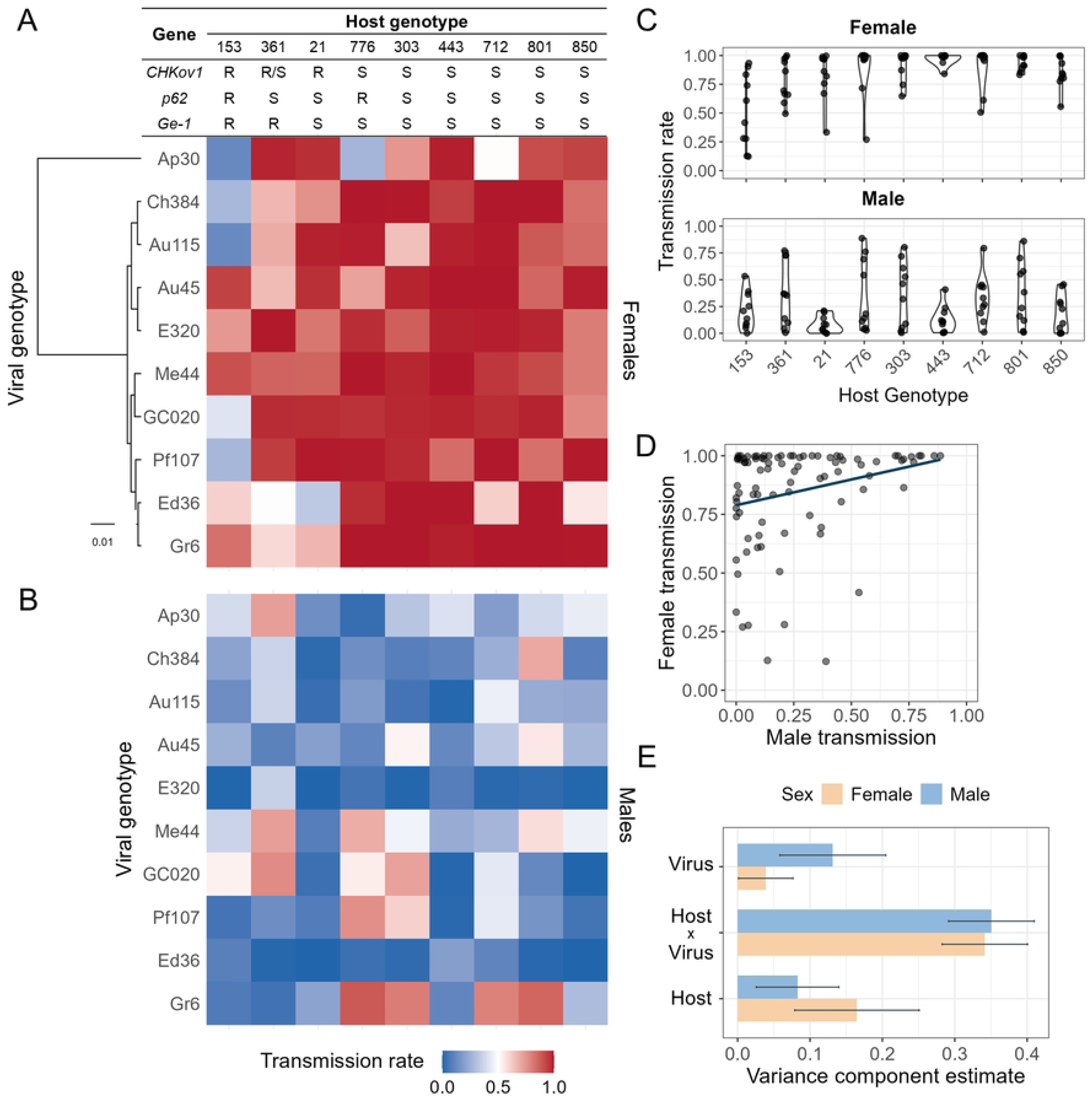
The effect of host and viral genotype on vertical transmission rate. (A) The proportion of progeny that were infected from an infected mother crossed to an uninfected father. Infection was detected using the CO_2_ assay. Host genotypes are annotated for three loci with known resistance (R) or susceptibility (S) alleles; heterozygotes are denoted R/S. The phylogeny of the virus strains is shown to the left; all nodes are supported with posterior probability ≥99.9%. (B) The proportion of progeny that were infected from an infected father crossed to an uninfected mother. (C) The distribution of transmission rates across host genotypes. (D) Phenotypic correlation between maternal and paternal transmission. Each point is a unique combination of host and viral genotypes. (E) Variance components estimated from an ASReml model with standard errors. The data supporting panels A–D can be found in S8 Data, and the values for panel E are provided in Table B in S1 Text.

In line with known transmission patterns, we observed strong sex differences: on average infected females transmitted the virus to 85% of their offspring, compared to just 26% for infected males (Fig 3C and 3D). Across all host–virus combinations, transmission rates were correlated between sexes (phenotypic correlation: *r_p_* = 0.5, *p* < 0.001; Fig 3D). However, while low-transmitting female genotypes never exhibited high male transmission, the reverse is not true (Fig 3D). The existence of lines where transmission by females is near-perfect but transmission through males is strongly reduced suggests specific barriers that may affect entry into the male germline. Despite this disparity in mean transmission, variability was similar between sexes (female standard deviation = 0.28; male standard deviation = 0.32).

In females, the combined genetic effects of the host or virus accounted for 55% of total transmission variance. However, this does not correspond to high genetic variation in viral fitness (as averaged across hosts), because the majority of this genetic variance in transmission was attributed to host–virus genotype-by-genotype (G_H_×G_P_) interactions (63%; Intraclass Correlation Coefficient: ICC = 0.34 ± 0.06 SE, *p* < 0.001; ICC represents proportion of total phenotypic variance; Fig 3E; all genetic variance components are shown in Table B in S1 Text), while only 7% was explained by genetics of the virus alone (ICC = 0.04 ± 0.04 SE, *p* = 0.076; Fig 3E). Therefore, a virus that has high transmission in one host genotype will frequently have low transmission in future generations when it finds itself in a different genotype. For instance, despite the virus strain Ap30 having high transmission in most host genotypes, transmission was strongly reduced in host lines 153 and 776. In this case this likely reflects sensitivity to the resistant allele of *p62* (Fig 3A).

The pattern was similar in males (Fig 3E). Genetic influences of the host or virus again explained the majority of transmission variance (56%). However, G_H_×G_P_ interactions contributed the largest proportion (62% of genetic variance; ICC = 0.35 ± 0.06, *p* < 0.001), exceeding the effect of viral genotype alone (main effect virus: 23% genetic variance; ICC = 0.13 ± 0.07 SE, *p* < 0.001). For example, virus strain Gr6 had the highest mean transmission in males (0.45), more than seven-fold higher than the lowest-transmitting strain (Fig 3B and 3C). However, this was driven by high rates in four host genotypes, with strongly reduced transmission in other lines.

For a vertically transmitted pathogen, transmission rate also affects host fitness. High transmission rates mean more of an individual’s offspring suffer the costs of infection. Mirroring the effects in viruses, G_H_×G_P_ interactions strongly reduced the influence of this factor on host fitness. In males, host genotype explained 15% of the genetic variance (ICC = 0.08 ± 0.06 SE, *p* = 0.006) compared to 56% explained by G_H_×G_P_ term (Fig 3E). In females, host genotype explained 30% of the transmission genetic variance in females (ICC = 0.17 ± 0.09 SE, *p* < 0.001), compared to 55% explained by G_H_×G_P_ interactions (Fig 3E). Therefore, as was the case in the virus, despite high levels of host genetic variation, the host fitness variation is greatly reduced by G_H_×G_P_ interactions.

Despite the phenotypic correlations between sexes (Fig 3D), the underlying genetic architecture of transmission was markedly sex specific. Notably, host–virus interaction effects, which dominated the genetic variance in both sexes, were only partially correlated (𝑟_𝑔_= 0.33 ± 0.12 SE; all genetic correlations are shown in Tables C and D in S1 Text), indicating substantial genotype-by-sex interactions. Viral genotype effects were moderately correlated across sexes (𝑟_𝑔_= 0.57 ± 0.39 SE), while the correlation of host genetic effects was poorly estimated (𝑟_𝑔_= 0.19 ± 0.46). These results suggest that while transmission phenotypes are correlated across sexes, there remain widespread sex-specific host–virus interactions.

### Sex specific interactions between host and virus genotypes determine viral load

Viral load is frequently a key determinant of host and viral fitness, mediating both transmission rates and virulence. We quantified viral load in 712 females and 722 males across all 90 host–virus genotype combinations (Fig 4A and 4B), observing a significant phenotypic correlation between sexes (*r_p_* = 0.5, *p* < 0.001; Fig 4D), indicative of shared genetic determinants.

**Fig 4.**
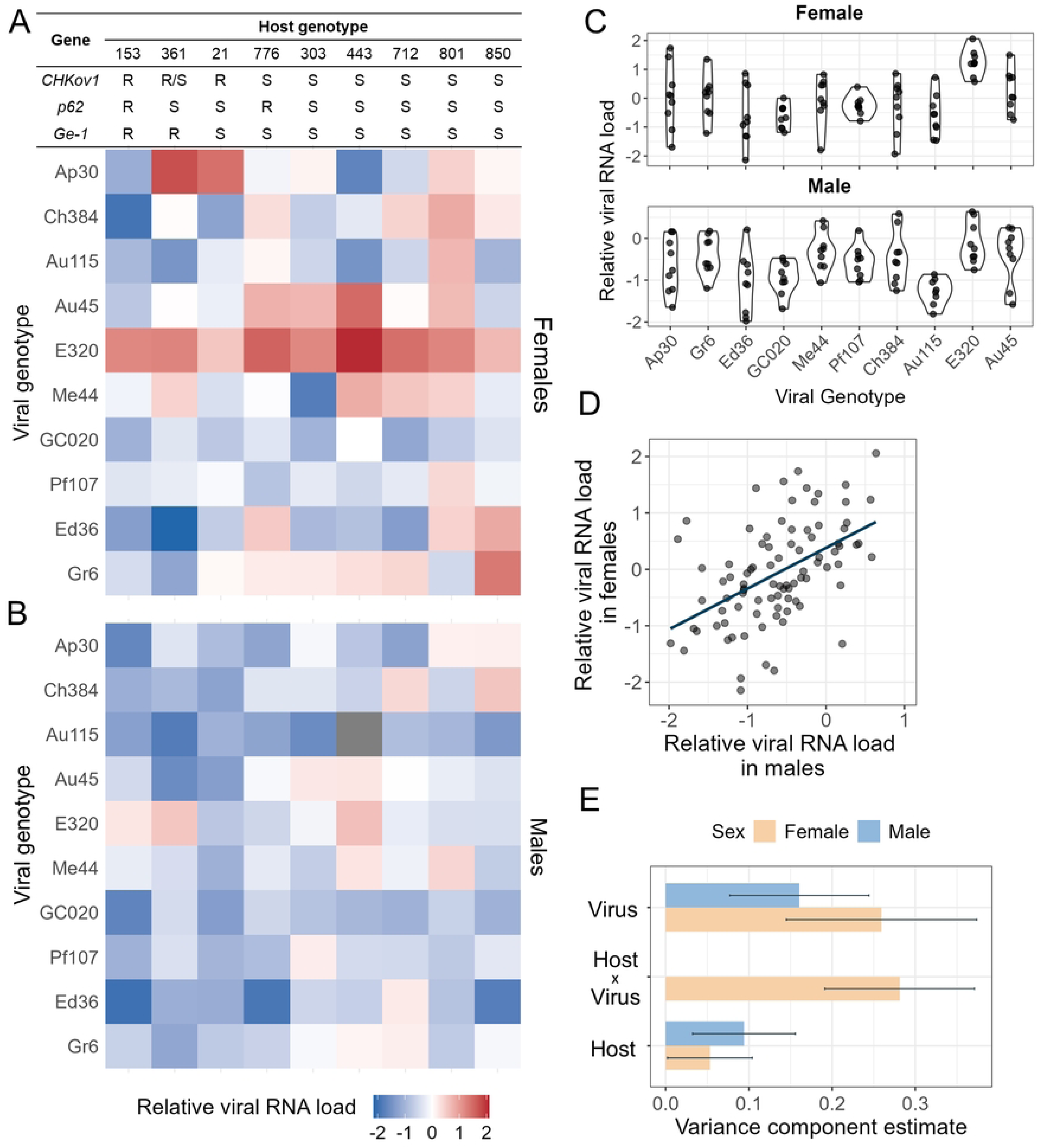
The effect of host and viral genotype on viral load. (A) Viral load in females across 90 combinations of *D. melanogaster* lines and sigma virus strains. (B) Viral load in males across the same host–virus combinations. Viral RNA was quantified by qPCR relative to the mean-centered expression of the host gene *RpL32*. Viral load estimates are based on 364 RNA samples from ∼1,630 flies. (C) Distribution of viral load by virus genotype. (D) Correlation of viral load between sexes across all unique host–virus combinations. Line indicates linear regression. (E) Variance component estimates from ASReml mixed models with standard errors. The absence of a male interaction term reflects a variance component not detected in the model. The data supporting panels A–D can be found in S8 Data, and the values for panel E are provided in Table B in S1 Text.

In females, viral load varied by 18-fold, with 59% of phenotypic variance attributable to genetic effects. Notably, G_H_×G_P_ interactions accounted for the largest share of the total genetic variance (47%; ICC = 0.28 ± 0.09 SE, *p* < 0.001; Fig 4E). For instance, virus strain Me44 typically exhibited high viral titres across hosts but showed markedly reduced load in line 303 (Fig 4A). Viral genotype alone explained 44% of the total genetic variance in females (ICC = 0.26 ± 0.11 SE, *p* < 0.001), with the highest-titre strain (E320) reaching more than four times the mean load of the lowest (GC020; Fig 4A and 4C). Although all three known host genetic polymorphisms that restrict sigma virus replication were present among the DGRP lines, their effects were inconsistent across viral genotypes (Fig 4A and 4C). Consequently, host genotype alone explained only 9% of the total genetic variance in females (ICC = 0.05 ± 0.05 SE, *p* = 0.067). Nonetheless, the line with the lowest overall viral load (line 153) was the only genotype carrying the resistant allele at all three loci (Fig 4A).

In males, genetic contributions to viral load were substantially reduced, with only 25% of phenotypic variance explained by host and viral genotypes. Unlike in females, G_H_×G_P_ interactions were not detectable (Fig 4E). Viral genotype accounted for the majority of total genetic variance in males (63%, ICC = 0.16 ± 0.08 SE, *p* < 0.001; Fig 4E), with strain E320 again exhibiting the highest viral load (Fig 4B and 4C). Host genotype explained the remaining 37% of total genetic variance (ICC = 0.09 ± 0.06 SE, *p* = 0.003; Fig 4E).

Together, these findings reveal sex-specific genetic architectures underlying viral load. G_H_×G_P_ interactions are a major contributor in females but are undetectable in males, suggesting fundamental sex differences in the genetic basis of infection outcome. In contrast both host and viral genetic effects on viral load were strongly correlated between sexes (main effect host genotype: 𝑟_𝑔_ = 0.71 ± 0.39 SE; main effect virus genotype: *r*_𝑔_ ≈ 1).

### Transmission is positively genetically correlated with viral load

The relationship between pathogen load and transmission potential is a common assumption in disease ecology [50]. While high pathogen loads are frequently associated with increased transmission rates in horizontally transmitted systems [44], in vertically transmitted parasites the relationship is more variable than this generalisation suggests, with both the strength and direction differing across host–parasite systems [62]. Although it has received theoretical attention [63], to our knowledge no study has separated the host and viral genetic contributions to this relationship empirically. We found that transmission is positively correlated with viral load in both females (*r_p_* = 0.35, *p* < 0.001; Fig 5A) and males (*r_p_* = 0.21, *p* = 0.044; Fig 5B), suggesting that physiological mechanisms linking replication and transmission—such as viral access to gametes—are at least partly shared across sexes, though more tightly coupled in females.

**Fig 5.**
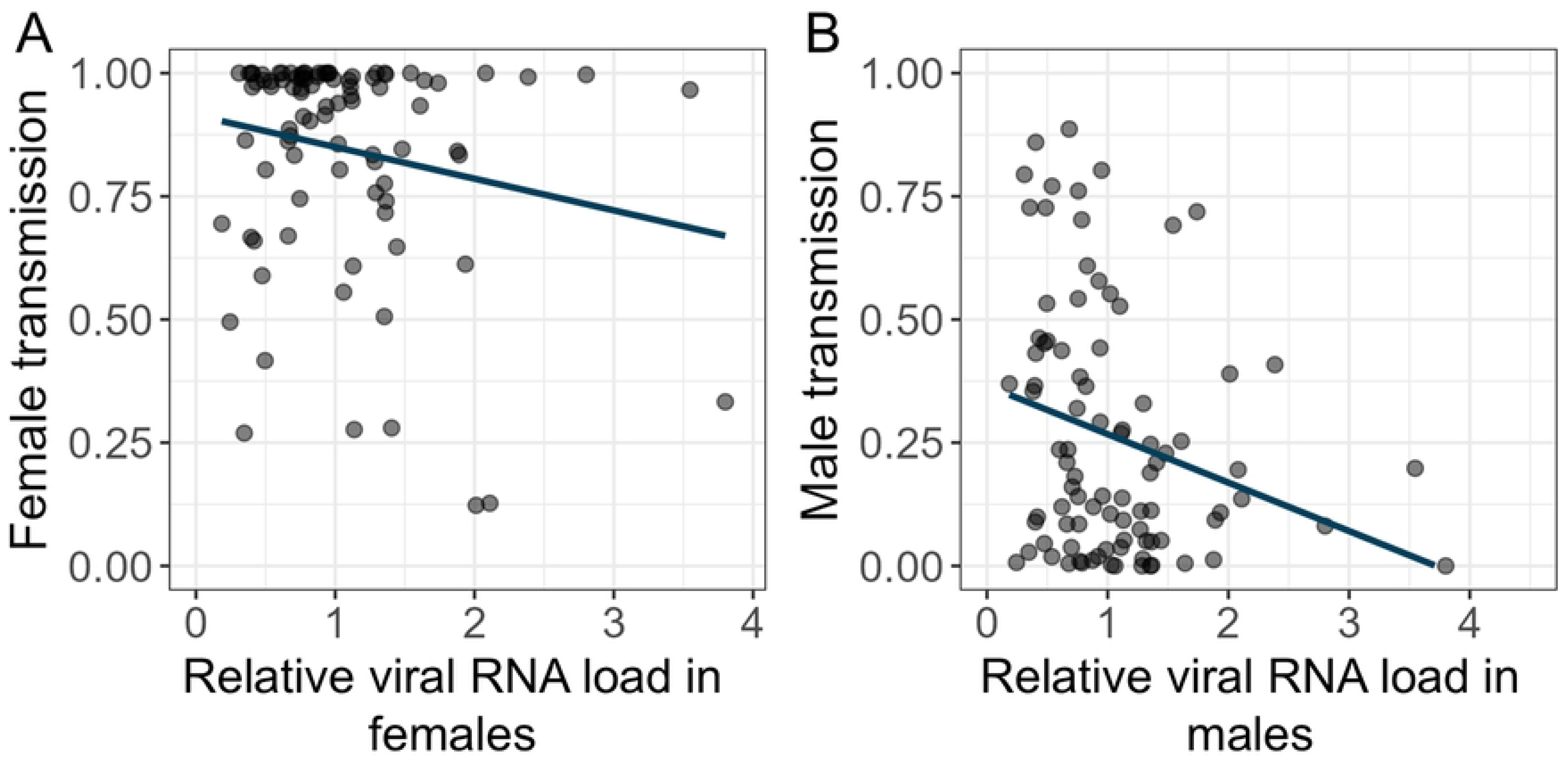
The impact of viral load on the transmission rate. The relationship between viral load and the proportion of progeny that were infected (A) maternally and (B) paternally. Viral load is relative to transcripts of the host gene *RpL32* and is the mean of males and females with the two sexes weighted equally. Each point is a unique combination of host and viral genotypes. Linear regression lines are shown. The data supporting these figures can be found in S8 Data.

To disentangle the genetic contributions to this correlation, we partitioned cross-trait genetic correlations (*r_g_*) by host genotype, viral genotype, and host–virus genotype interactions. Although most 𝑟_𝑔_ estimates exhibited substantial standard errors, consistent trends emerged. In females, virus genetic effects (𝑟_𝑔_≈ 1), host-virus interaction effects (𝑟_𝑔_= 0.42 ± 0.15 SE), and host genetic effects (𝑟_𝑔_ = 0.46 ± 0.34 SE) all showed positive correlations between viral load and transmission. In males, host and virus genetic effects on viral load and transmission were positively correlated (host: 𝑟_𝑔_= 0.52 ± 0.42 SE; virus: 𝑟_𝑔_= 0.32 ± 0.39 SE), while interaction effects were undetectable (Table C in S1 Text). This suggests genetic factors determining transmission are partially mediated by viral load, but other processes also play a key role.

### Genetic interactions between host and virus reduce the effect of virulence on host fitness

The fitness consequences of infection for the host are determined by viral virulence—the degree of harm inflicted on the host. Because vertically transmitted viruses depend on host survival and reproduction for their own propagation, virulence also directly constrains viral fitness in this system. We quantified virulence across 90 host–virus genotype combinations by measuring fecundity (eggs laid) and hatch rate (proportion of eggs hatched; Fig 6A and 6B). After normalizing for baseline differences among host genotypes, we found hatch rate and fecundity were positively correlated (*r_p_* = 0.48, *p* < 0.001; Fig 6C), suggesting both reflect overall reproductive success.

**Fig 6.**
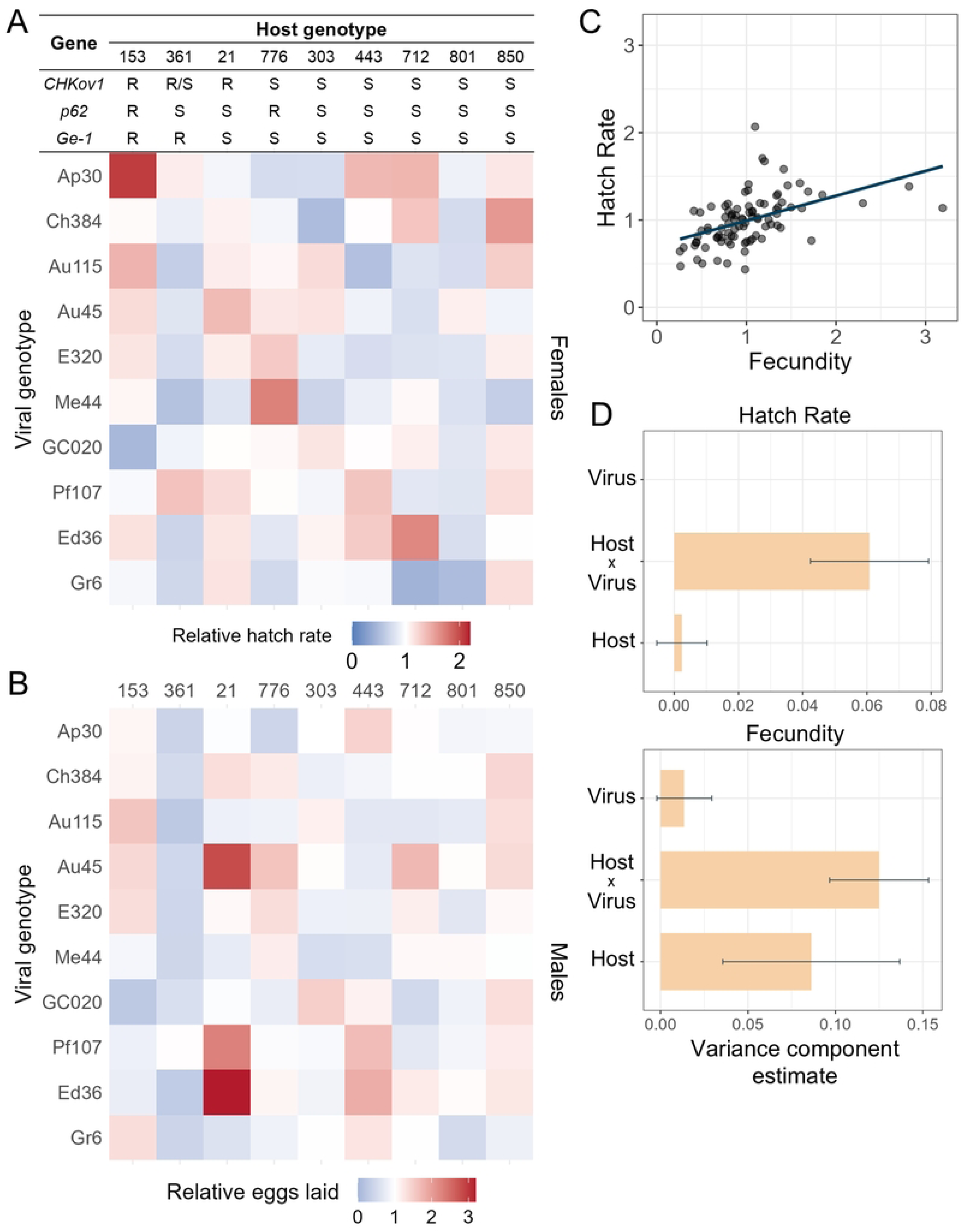
The effect of infection on female reproductive success. The effect of viral infection on (A) The hatch rate and (B) number of larval offspring. Both statistics are relative to the estimate from uninfected flies. (C) The correlation between the number of eggs laid and the proportion of eggs that hatch. Each point is a unique combination of host and viral genotypes. Eggs were laid by individual 6-7 day-old females over a 24-hour period, and the proportion that hatched counted 48 hours later. (D) The variance in hatched eggs explained by the host and virus genotypes from an ASReml model. Error bars indicating standard errors. The absence of a male interaction term reflects a variance component not detected in the model. The data supporting panels A–D can be found in S8 Data, and the values for panel E are provided in Table B in S1 Text.

To partition genetic effects, we included mean trait values of uninfected lines as covariates to estimate the impact of infection on fecundity. We found strong positive genetic correlations between hatch rate and fecundity explained by host genetics (𝑟_𝑔_ = 0.97 ± 0.34 SE) and host-parasite interaction (𝑟_𝑔_ = 0.97 ± 0.11 SE). However, the combined effects of host and viral genetics accounted for only 6% of the phenotypic variance in hatch rate and 10% in fecundity—substantially less than for viral load or transmission—indicating high residual variability in reproductive traits. Within the genetic component, nearly all of total genetic variance in hatch rate was attributable to G_H_×G_P_ interactions (96%, ICC = 0.061 ± 0.018 SE, *p* < 0.001; Fig 6D). G_H_×G_P_ effects also explained 56% of the total genetic variance in fecundity (ICC = 0.125 ± 0.028 SE, *p* < 0.001; Fig 6D).

The main effect of viral genotype did not significantly influence hatch rate or fecundity (Fig 6A and 6D), indicating no detectable impact of virulence on viral fitness at the population level. Host genotype effects were similarly weak for hatch rate (ICC = 0.002 ± 0.008 SE, *p* = 0.37) but accounted for 38% of the total genetic variance in fecundity (ICC = 0.086 ± 0.051 SE, *p* < 0.001, Fig 6A and 6B), possibly reflecting differences in host tolerance to infection.

We next tested for a trade-off between transmission and virulence, expected if higher replication enhances transmission at the expense of host fitness [44,64]. Phenotypically, transmission rate was negatively correlated with both fecundity and hatch rate (Fig 7A, 7B and S5 Fig), consistent with such a trade-off. Despite these phenotypic patterns, neither transmission nor viral load showed a significant genetic correlation with fly fitness (Table C and D in S1 Text).

**Fig 7.**
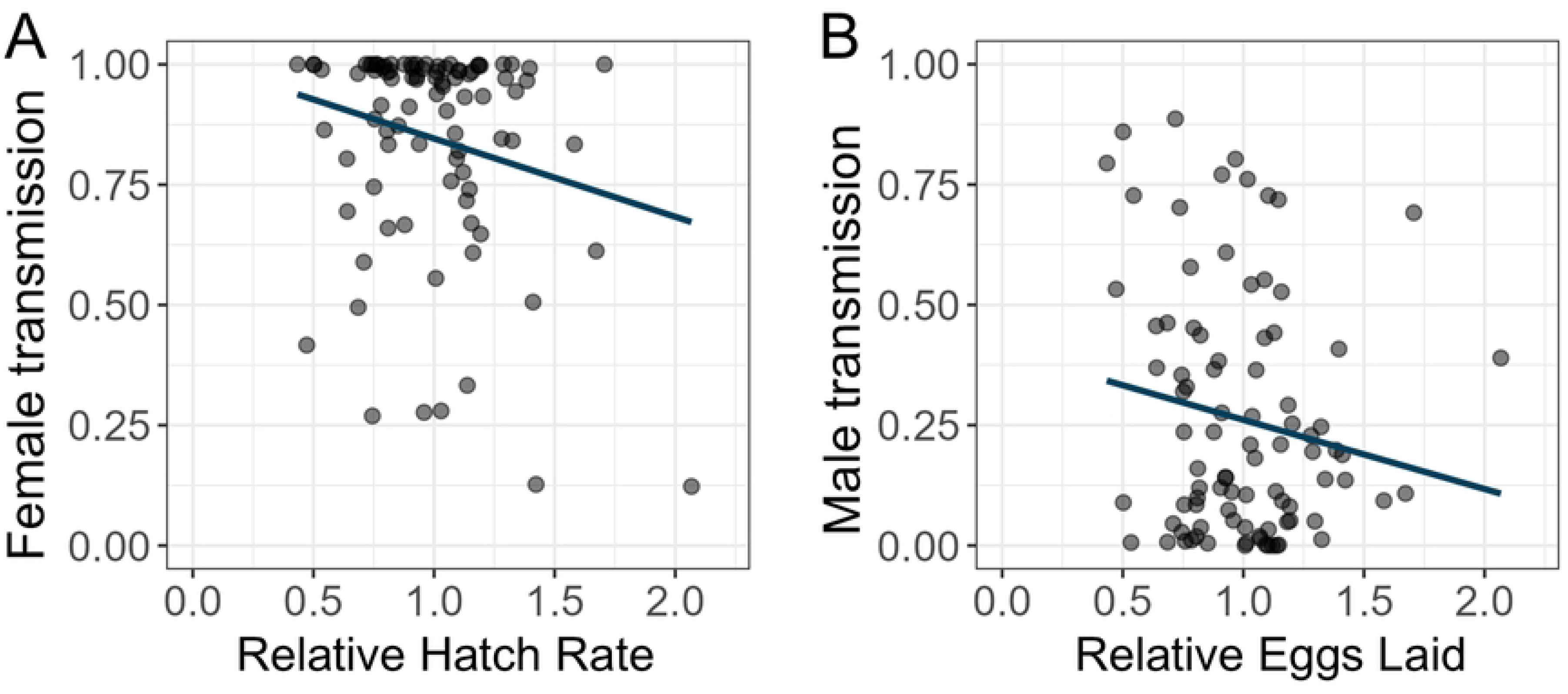
Correlation between fitness traits and viral transmission. (A) Correlation between the number of eggs that hatched and transmission in females. (B) Correlation between the number of laid eggs and transmission in males. Each point is a unique combination of host and viral genotypes. Linear regression lines and equations are shown. The data supporting these figures can be found in S8 Data.

## Discussion

A central question in coevolution is what maintains genetic variation in interacting partners despite ongoing selection. One explanation lies in genotype-by-genotype effects on fitness that can in turn enable negative frequency-dependent selection, in which no genotype gains a permanent advantage. This shapes the structure of fitness variance: the fitness of any genotype depends on which partner genotypes it encounters, shifting genetic variance from additive main effects into the interaction term. This will reduce the additive genetic variance in fitness and slow rates of adaptation. To investigate how strong this pattern is in a natural system, and what it reveals about coevolutionary dynamics, we investigated the *Drosophila*–sigma virus system, combining theoretical simulations with large-scale experimental tests.

To establish theoretical expectations, we first modelled host-parasite coevolution in a system with vertical transmission. The dynamics are highly sensitive to parameter choice. Under some parameter values, coevolution produced damped oscillations converging on a stable equilibrium. Pure stable equilibria reached without oscillation were rare. Other parameters resulted in persistent or unstable oscillations. Across these regimes, fitness variance for both host and virus were dominated by the *V_G×G_* interaction term which contained essentially all fitness variance at equilibrium and typically remained the major component through oscillations even as additive genetic variance was transiently exposed. This pattern is robust across both gene-for-gene and matching-allele interaction models, indicating it is a general consequence of host-parasite coevolution with vertical transmission. The parameter space producing these dynamics is consistent with infection parameters measured in natural *Drosophila*–sigma virus populations [24,29–33]. Thus, across much of the biologically plausible parameter space, a snapshot of a natural population is expected to show *V_G×G_* dominating and additive genetic variance in fitness eroded.

Next, we carried out a large-scale study with more than 4000 flies across 90 combinations of host and virus genotypes to quantify G_H_xG_P_ interactions between *Drosophila melanogaster* and sigma virus. We measured key components of host and viral fitness—transmission, fecundity and egg hatch rate— and viral load, a potential mediator of fitness. For all these parameters we estimated the contributions of host genetics, virus genetics, and their interactions to their variance.

Our results consistently show that infection outcomes depend on specific G_H_xG_P_ interactions across all measured traits: no host genotype was universally resistant, and no viral genotype was universally infective. Previous work in this system showed that a *p62* resistance allele affected maternal transmission of particular viral isolates and had no effects on isolates that had evolved to escape the effects of this allele [37,39,65,66]. On the other hand, Wilfert & Jiggins found that with high genetic variation in paternal transmission across both host and viral genotypes, the host resistance gene remained effective across virus isolates, suggesting predominantly general resistance [40]. Carpenter et al. similarly found positive genetic correlations between transmission rates across host genotypes, though the fact these correlations were below one indicated that there was a significant G_H_xG_P_ component [66]. We find G_H_xG_P_ dominating the genetic variance across every fitness-related trait we measured. Fitness in this system is therefore largely a property of specific host–parasite combinations, with little additive genetic variance available for selection on either party alone.

While the relative strength of G_H_xG_P_ interactions varies across host-parasite systems [12,19,67–73], our study revealed strong G_H_xG_P_ interactions governing most sigma virus infection outcomes. Therefore, despite extensive genetic variation in both host and sigma virus, there may be substantially less variation in the mean fitness of either party. In this case interactions among genotypes maintain coexistence without any genotype becoming universally dominant [74]. At a stable equilibrium the variance explained by G_H_xG_P_ interactions is maximized, while host and parasite individual genetic variance approaches zero. This interaction variance is in effect cryptic: inaccessible to directional selection under the current distribution of partner genotypes but exposed when that distribution shifts. In practice we expect natural populations to spend most of their time away from any stable equilibrium, because mutation, environmental change, and demographic effects shift the underlying parameters faster than the system relaxes. Such perturbations increase the statistical weight of host or pathogen main effects, and release previously hidden variance, potentially enabling rapid adaptive change. In parameter space where genotypes cycle, shifts in allele frequency of the opposing party has the same effect, periodically shifting the variance to the main effect, allowing rapid adaptation.

Consistent with prior research [5,24], we observed significant sex differences in transmission rates. Infected females transmitted the virus to 85% of their offspring on average, compared to 26% for infected males. While maternal transmission is the primary route of sigma virus spread, male transmission is essential for viral persistence at the population level: in its absence, the fitness costs of infection would gradually reduce the frequency of infected females, reducing viral prevalence [51,61]. For transmission rates, summed genetic effects explain over half the variance in both sexes, with similar magnitudes in males and females. Therefore, there are comparable influences of maternal and paternal transmission on fitness variance, and there is expected to be antagonistic selection on *Drosophila* to reduce transmission to offspring and the virus to increase transmission.

For fly life history traits, genetic effects explain far less variance than was the case for viral load or transmission. Nevertheless, G_H_xG_P_ were substantial, with host genetics playing a prominent role in fecundity. Intriguingly, despite well-documented fitness costs of sigma virus infection [27,28,33,61,75,76], our results indicate only mild effects on female reproduction. Moreover, infection frequently enhanced this trait, mirroring weakly supported reports of minor reproductive gains in infected males [59,77]. The reasons for these reproductive increases and whether they are relevant in the field remain unclear.

Unlike transmission, viral load showed a strong phenotypic correlation and relatively high genetic correlation between males and females, mediated by host and parasite genetic effects. Therefore, host and parasite genetics exert shared control over viral replication in both sexes. In females, there is a high genetic correlation between viral load and transmission, consistent with prior observations tying elevated ovarian viral load to transmission rates [30,45]. In males, this correlation is weaker, and high viral load does not reliably predict paternal transmission success. This discrepancy likely arises because male transmission requires the virus to infect sperm cells, potentially influenced by host genetic factors that prevent viral access to the germline [5,30,78,79]. Males also show a broader range of infection-induced gene expression changes compared to females, possibly reflecting specific resistance responses in the male germline [80]. The weak genetic correlation between male and female transmission rates further supports sex-specific barriers to transmission [5,80].

We also examined correlations between other infection-related traits. These estimates are best viewed as preliminary biological trends rather than conclusive findings as genetic correlations were weak, with most not reaching statistical significance. Transmission rate showed negative phenotypic correlations with both fecundity and hatch rate, suggesting a reproductive cost of higher transmission. Although viral load robustly affected transmission, it did not have a robustly detectable effect on fly fitness. Therefore, we cannot conclude that viral replication underpins a trade-off between virulence and transmission.

The generally weak genetic correlations among infection-related traits suggest the G_H_xG_P_ interactions often have trait-specific effects. This trait-specificity, as observed in *Drosophila* anti-parasitoid responses, argues against the same genetic loci governing different aspects of host-virus interactions [81]. Instead, several independent genetic factors likely associated with different traits shaping outcomes [82,83].

We did not aim to dissect the roles of known *Drosophila* resistance genes or sigma infectivity architecture, so we could not isolate individual resistance allele effects. However, our fly lines carried both resistant and susceptible alleles of three major-effect genes (*p62*, *CHKov1*, *Ge-1*). While major-effect genes have traditionally been emphasized in studies of host resistance [15], our data suggest a complex architecture, as phenotypic variation for all traits was similar between fly groups with and without major resistance alleles.

Host–parasite coevolution has long been proposed to maintain genetic diversity through genotype-specific fitness effects, yet empirical demonstrations of how these interactions structure variance across different components of fitness have remained rare. Our results show that, in a natural vertically transmitted virus system, genetic variance across a suite of host and virus traits was dominated by the *V_G×G_* interaction term, consistent with coevolution masking heritable variation from the effects of natural selection. This observation aligns with theoretical predictions that negative frequency-dependent selection shifts fitness effects into interaction components, thereby slowing adaptation and promoting coexistence. Furthermore, G_H_×G_P_ interactions were frequently sex- and trait-specific, which may further reduce the additive genetic variance in fitness. Comparable patterns of strong genotype-by-genotype effects have been reported in diverse systems, yet the magnitude and consistency of these effects across multiple fitness-related traits in our study suggest that such interactions may be more pervasive than previously appreciated. These results highlight that coevolution can generate extensive cryptic genetic variation, which may be revealed either when ecological or evolutionary conditions shift, or when the process of coevolution itself alters the direction of selection. Our results illustrate the importance of considering genotype-specific interactions when predicting evolutionary responses to emerging pathogens, environmental change, or interventions that alter the distribution of host or parasite genotypes.

## Materials and Methods

### Simulation model

We implemented a model to examine the dynamics of coevolution and how fitness variance is partitioned between additive and interaction components. The basic version of the model, a single pathogen with no genetic variation in either host or parasite, is shown schematically in Fig 8 and specified below. The full model extends this to two co-circulating viral genotypes and two biallelic host resistance loci that coevolve. Simulations were implemented in R (v.4.4.0) in Rstudio. The complete derivation of the full model, including the interaction matrices for each of the coevolution regimes considered here, is given in S1 Text.

**Fig 8.**
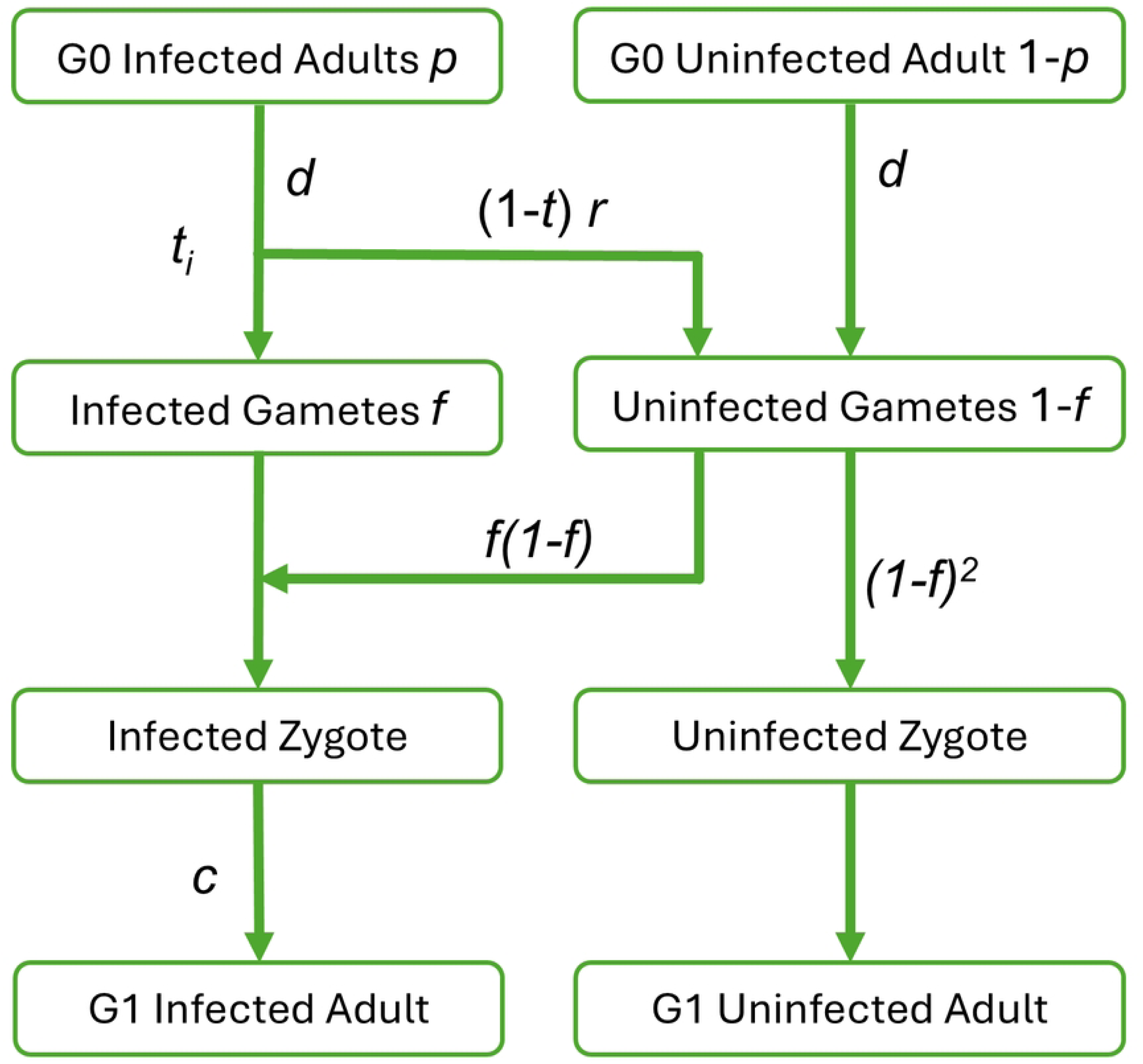
Schematic of the model of a vertically transmitted pathogen. The schematic shows the basic single-pathogen, single-locus version with no genetic variation, corresponding to equations (1)–(3); the full model incorporates two pathogen genotypes and two host resistance loci. Parameters: *t*, pathogen transmission rate (in the full model *t*ᵢ for virus strain *i*, further reduced to *t_ᵢ,G_* = *t*ᵢ(1 − *r* · 1*_ᵢ,G_*) in hosts carrying a matched resistance allele); *c*, cost of infection on offspring viability (a single parameter, applied equally to hosts infected with either virus); *r*, reduction in pathogen transmission in hosts carrying a matched dominant resistance allele (dominant); *d*, fertility cost per locus at which the host carries a dominant resistance allele (dominant). In the simple model *r* and *d* act uniformly on every host; both become dynamically relevant only when resistance varies across host genotypes in the full model.

The model tracks cohorts of infected (frequency *p*) and uninfected (frequency 1 − *p*) adult hosts across discrete, non-overlapping generations. Each adult produces gametes and hosts that carry a matched dominant resistance allele incur a fertility cost *d*, so each such adult’s per-capita gamete output is (1 − *d*). Each infected parent transmits the pathogen to a fraction *t* of its gametes, with equivalent transmission through males and females; the pathogen’s transmission is further reduced by a factor *r* in hosts that carry the resistance allele. The frequency of infected gametes (p̅) is the ratio of infected to total gametes in the population:

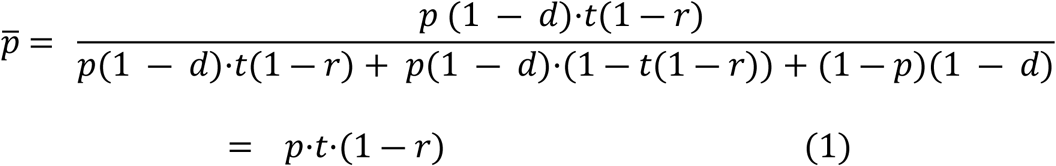

In the simple model *d* acts uniformly on every adult and so factors out of every term in the gamete pool, leaving *p̅* to depend only on *p*, *t*, and *r*.

Random union of male and female gametes yields zygotes. A zygote is infected either if the maternal gamete carries the pathogen, or if the maternal gamete is uninfected and the paternal gamete is infected; the resulting infected zygote frequency *f* is

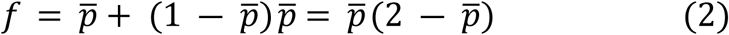

Infected zygotes develop at a relative viability cost *c*. After applying the viability cost, the frequency of infected adults in the next generation is

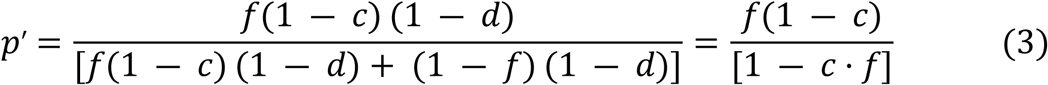

Both, *r* and *d* parameters therefore have no dynamical effect in the simple model, as they drive coevolutionary dynamics only once they vary across host genotypes in the full model (S1 Text).

The full model extends equations (1)–(3) by letting resistance and transmission depend on genotype. The infection cost *c* is retained as a single parameter applied equally to hosts infected with either virus, but each viral strain *i* has its own base transmission rate *tᵢ* (*t₁* and *t₂* set independently), and *tᵢ* is further reduced in hosts carrying a matched resistance allele, giving the effective transmission rate *t_ᵢ,G_* = *tᵢ*(1 − *r · 1_ᵢ,G_*), where 1*_ᵢ,G_* is a resistance indicator (1 if host *G* is resistant to virus *i*, 0 otherwise). The fertility cost *d* is then applied at the gamete stage, multiplicatively per locus at which the host carries a dominant resistance allele, so host genotypes now differ in their gamete output (a host resistant at both loci pays (1 − *d*)²). The difference *t₁ − t₂* represents an infectivity cost of escape from host resistance when *t₂ < t₁* — the parasite-side counterpart of the host’s fertility cost *d*.

At each generation, fitness variance across the joint host–virus genotype distribution is decomposed into host-genotype, virus-genotype, and host × virus interaction components, for both host and virus fitness (S1 Text).

The simulations track the frequencies of host genotypes and their infection status across generations. The full model maintains a two-locus structure with two biallelic resistance loci (*A/a* and *B/b*) and two viral genotypes, coevolving the nine possible diploid host genotypes with the two viruses. Three coevolution models were implemented. In the single-locus gene-for-gene models (GFG1), shown in the main text, dominant allele *A* confers resistance to Virus 1 and gene B is absent. In the two locus gene-for-gene model (GFG2), a dominant allele *B* of a second unlinked host gene confers resistance to Virus 2. In the matching-allele model (MA), allele *A* confers resistance to Virus 1 while the susceptible genotype (*aa*) confers resistance to Virus 2 — the same locus determines resistance to both viruses, in opposite directions depending on genotype. A selective sweep model was additionally implemented. This is identical to GFG2, but with no costs associated with resistance or the virus overcoming resistance, and the sequential introduction of Virus 2 and allele *B* at fixed generations, simulating directional arms-race dynamics. Full model mechanics and interaction matrices are provided in S1 Text.

### Parameter sweep

To characterize the parameter space supporting different coevolutionary outcomes, we systematically varied key parameters of the GFG1 model across biologically plausible ranges informed by field estimates from natural *Drosophila*–sigma virus populations, with extended ranges to explore model behavior outside the empirically observed window. For each parameter combination we recorded mean resistance-allele and virus-genotype frequencies, the fitness variance components *V_G(host)_*, *V_G(virus)_,* and *V_G×G_*, and the qualitative type of coevolutionary dynamics. Full details of the parameter-sweep procedure are given in S1 Text. The R script used to run all coevolutionary simulations is available on Zenodo: [DOI: 10.5281/zenodo.20083947].

### Fly lines

We used nine *D. melanogaster* lines from *Drosophila* Genetic Reference Panel (DGRP) [60], six of which were kindly provided by Pedro Vale, University of Edinburgh. These lines were selected based on the fitness scores [84] and variants of the genes known to confer resistance to DMelSV (*Ge-*, *CHKov-1*, *p62* (*ref2p*) [35–37,65,85]. Line 153 carries resistance alleles at all three genes, line 361 carries resistance alleles at *Ge1* and heterozygous for *CHKov-1*, line 21 carries resistance alleles at *CHKov-1* only and line 776 carries resistance alleles at *p62* only, lines 303, 443, 712, 801 and 850 are susceptible at all genes. We confirmed the alleles by PCR with primers targeting susceptible and resistant alleles [15].

### Virus isolates

Ten sigma virus isolates from geographically diverse *D. melanogaster* populations were used, including publicly available Ap30 - United States; Florida, PF107 – Athens, Greece; GC020 - Galicia, Spain; E320 - Essex, UK [40,86]; and strains collected from the wild fly populations during summer 2021 in the United Kingdom: Me44 - Cambridgeshire; Ed36 - Edinburgh; Gr6 - Sussex; Au45 and Au115 - Perth and Kinross; and Ch384 – Cambridge. To identify unique genotypes of the viral isolates from the same collection locations and to confirm the correct genotypes of the publicly available isolates we sequenced part of the viral genome covering the region from the gene P to the beginning of the gene L region by Sanger sequencing. Sequences were aligned using MUSCLE and then trimmed by eye to remove regions with missing data [87]. Phylogenetic reconstruction was performed using BEAST v1.10.5.0 with 6332bp of sequences fitted to an uncorrelated relaxed lognormal molecular clock model using a speciation birth-death process tree-shape prior [88]. A HKY substitution model with a gamma distribution of rate variation with four categories and a proportion of invariable sites was used. Models were run for 100,000,000 steps using a 10% burn-in. Sequence alignment was also used for designing qRT-PCR primers (Table E in S1 Text).

### Generation of infected lines

Infected host–virus lines were established by pairing each of the nine host lines with each of the ten viral isolates. Viral isolates were introduced into DGRP host genetic backgrounds using a backcross scheme adapted from Wilfert & Jiggins (2010) [40], using the translocation balancer stock *T(2;3)*CyO-TM8 (CyO, *Adh*[nB]: TM8, *l(3)DTS4*[1], Bloomington). For published viral isolates, infection was first established in *w*^1118^ (DrosDel)[89] adults by intra-abdominal injection of crude homogenates prepared from infected fly tissue, following standard sigma virus injection protocols[90]. Once infection was established, infected *w*^1118^ females were crossed to males of the balancer stock (Generation 1 cross: *w^1118^* ♀ × *T(2;3)*CyO-TM8 ♂). Infected F1 females carrying *T(2;3)*CyO-TM8 were crossed to males of the target DGRP line (Generation 2 cross: F1 infected *T(2;3)*CyO-TM8 / *w^1118^* ♀ × DGRP ♂). From F2 offspring, infected siblings carrying the balancer chromosome were crossed to establish stabilised infected stocks (Generation 3 cross: F2 infected *T(2;3)*CyO-TM8 / DGRP ♀ × F2 infected *T(2;3)*CyO-TM8 / DGRP ♂). Stocks were then continued by selecting infected offspring not carrying the balancer chromosome. Established stocks were maintained for two or more generations at stable infection transmission rates before phenotypic assays were performed. After two generations of backcrossing, infected lines were considered to carry the autosomal genetic background of the recipient DGRP line. For wild-caught viral isolates, infected females collected from natural populations were crossed directly to males of the balancer stock as the Generation 1 cross (infected wild-caught ♀ × *w^1118^*; *T(2;3)*CyO-TM8 ♂). All subsequent crosses proceeded identically to those for the infected *w^1118^* isolates. Sigma virus infection status was confirmed at each generation selection step by CO_2_ sensitivity assay, in which actively infected flies exhibit prolonged paralysis upon exposure to CO_2_ gas [31].

### Experimental design

All assays were conducted using a randomised block design. Given the scale of the experiment (90 infected lines plus uninfected controls), assays were completed across several rounds with 10–30 lines per round. Approximately one-quarter of lines from each previous round were carried forward as internal replicates to allow correction for between-round variation. The number of DGRP lines per round was restricted to 2–3 to minimise the number of uninfected controls required. We maintained fly stocks and conducted all phenotypic tests at 25°C with 65% humidity and a 12:12h light:dark photoperiod. For all phenotypic tests, to stabilise sigma virus infection (transmission rates are higher through offspring infected by their mothers), infected parental flies used in tests were the F1 offspring of one infected female crossed to 2–3 males of the corresponding uninfected DGRP line.

### Viral transmission test

We tested the transmission rate of the different virus isolates in each host genetic background. Both the transmission and virulence tests across all 90 infected lines were performed in several rounds of crosses using a randomized block design, with 10–20 lines per round. Each round included approximately one-quarter of the infected lines from the previous round—specifically those with insufficient replicates. For the transmission test, we used individual virgin 2–3-day-old infected parental females or males from each of the 90 infected lines and a single uninfected fly of the opposite sex of the same age. Flies were kept in vials with cornmeal medium [15] supplemented with dried yeast for four days for egg laying. Next, 6–7-day-old parents were collected and tested for infection and immediately homogenised in 180µL TRIzol (Invitrogen, USA) and stored at −80°C for subsequent RNA extraction. The total number of the offspring and the number of infected flies were counted for each of the infected parents. A minimum of two biological replicates were used for grandparental flies and three for the parental flies for each infected line and sex tested.

### Virulence test

The impact of sigma infection on female reproductive performance was assessed by measuring changes in fecundity (number of eggs laid) and hatch rate (proportion of eggs hatched) in both infected and uninfected flies. The virulence test across all 90 infected lines was conducted following the same randomized block design as described above. For the control, we used uninfected DGRP lines, and all tests were performed in the same way as for the infected flies. For the tests, individual infected virgin females (parental test flies) were crossed with an uninfected male of the same age from the corresponding DGRP lines. For most lines, a minimum of two biological replicates for progenitor flies and two for parent flies were used for each infected line.

Flies were placed on yeasted cornmeal medium, then after four days transferred to vials containing apple-juice agar (2% agar) with 30 µL of yeast solution (1 mg live yeast in 3 mL H_2_O). After two further days, flies were transferred to vials with fresh apple-agar medium containing 0.5% activated charcoal (to provide contrast for counting) and 3 µL of yeast solution (1 mg dry yeast in 10 mL H_2_O) for 24 hours. The surface of the medium was brushed to create an optimal surface for oviposition. The number of eggs laid per vial was counted for infected and control flies. For the fertility test, 2 days after the fecundity test, hatched eggs were counted. Hatch rate was calculated as hatched eggs divided by total eggs laid. To account for variation in baseline fitness across DGRP lines, relative fecundity and relative hatch rate were calculated by dividing each infected-fly value by the mean for control flies from the same round.

### Measurement of viral RNA load

We measured the viral load in 6–7-day-old adult flies used in the transmission test for all combinations of viral and host genotypes and both sexes. For each infected fly line and sex, up to 14 randomly selected individual flies, homogenized in TRIzol, derived from up to six grandparental flies (see viral transmission test), were proportionally pooled in one sample for viral RNA extraction. RNA was extracted following the standard TRIzol extraction protocol.

In total, two pooled RNA samples were obtained for most infected lines and both sexes. We reverse transcribed RNA with GoScript reverse transcriptase (Promega, USA) and random hexamer primers. Viral RNA was quantified by qRT-PCR relative to the endogenous control housekeeping gene RpL32 in two technical replicates, using primers from Longdon et al. [91]. The viral RNA was amplified using primers targeting sequence common for all virus genotypes overlapping viral genes *X-M* (Table E in S1 Text). We performed qRT-PCR using cDNA diluted 1:10 with nuclease free water on the Applied Biosystems StepOnePlus system. The reaction was done with SensiFAST SYBR Hi-ROX (Meridian Bioscience, USA) and the following PCR cycle: 50°C for 2 minutes followed by 95 °C for 10 minutes and followed by 40 cycles of: 95 °C for 15 s followed by 60 °C for 60 s. Each technical replicate for either housekeeping or viral primers for all samples was placed on one 384-well plate. Samples where the Ct difference between technical replicates exceeded 1.5 for either gene were removed.

A large sex-specific difference in RpL32 expression was detected, requiring sex-specific normalization. Mean-centered RpL32 Ct values were calculated separately for females and males across biological replicates, then the overall mean RpL32 Ct was added back to ensure all values were positive and to make values comparable with the sigma Ct values. The normalized ΔCt (representing relative viral RNA load) was calculated as the difference between the mean-centered RpL32 Ct and the cycle threshold of the targeted sigma virus gene. Relative viral RNA load corresponds to log_2_ viral load.

### Statistical analysis

We used linear mixed models, fitted with ASReml-R to estimate genetic variance components for viral load, transmission rate, fecundity and hatching rate [92]. Given strong evidence of sexual dimorphism in viral load and transmission we modelled these as sex-specific traits in order to detect any genotype×sex interactions present. First, we fitted a univariate model to each of the 6 traits, with no fixed effects specified but random effects of the host and virus identities included, as well as a factor defined by the pairwise combination. This yielded estimates of the among-host genetic variance, the among virus genetic variance, and variance attributable to host-virus G_H_xG_P_ interactions, which we scaled to intra-class correlations (i.e. proportions of total phenotypic variance). For each trait, we tested the significance of the three genetic variance components using likelihood ratio tests (assuming a 50:50 mix of Χ^2^ distributions with 0 and 1 df following [93]. All models assumed Gaussian error structures. Variance components were constrained to be positive and in some instances estimates were bound to the edge of allowable parameter space. In such cases we interpret the point estimate variance as zero (with no standard error estimable).

Next, we formulated a multivariate mixed model to estimate among-trait genetic correlations at host, virus and host×virus levels. Fixed and random effects were as described for the univariate models. Random effect structures were modelled using generalised heterogeneous correlation matrices in ASReml-R, while a diagonal matrix (i.e. trait-specific variances but no covariances) was used for the residual structure. This is because different traits were measured on different flies (so observation-level residual covariance is not estimable). Traits were scaled to standard deviation units to facilitate model fitting, but despite this we could not obtain a stable convergence for the full (six-trait model). Consequently, results are from a five-trait model after dropping hatch rate. Genetic correlations with hatch rate and the other traits were then estimated using a series of bivariate models. The phenotypic correlations among virulence, transmission, and viral load were estimated by Pearson’s product-moment correlation.

## Data Availability Statement

All data necessary to replicate the findings of this study are provided as Supporting Information. Raw experimental data (viral load measurements, transmission rates, and reproductive fitness assays) are provided in S8 Data. The Sigma virus genomic sequences generated in this study are in GenBank [awaiting accession numbers].

## Supporting information

**S1 Text. Supplementary methods and statistical results.** Full description of the coevolutionary simulation model. Includes Table B in S1 Text. Genetic variance components (intra-class correlations) from univariate models; Table C in S1 Text. Genetic correlations from the multivariate (five-trait) model; Table D in S1 Text. Genetic correlations from bivariate models; and Table E in S1 Text. Primers used in this study.

**S2 Fig. Coevolutionary dynamics under a matching-allele model (MA).** Time series of allele frequencies and genetic variance components under damped (left) and persistent (right) oscillation regimes. The host carries one biallelic resistance locus where allele *A* confers resistance to Virus 1 and homozygous recessive aa confers resistance to Virus 2.

**S3 Fig. Coevolutionary dynamics under a two-locus gene-for-gene model (GFG2).** Time series of allele frequencies and genetic variance components under damped (left) and persistent (right) oscillation regimes. The host carries two unlinked biallelic resistance loci (*A/a* and *B/b*); allele *A* confers resistance to Virus 1 and allele *B* to Virus 2.

**S4 Fig. Selective sweep dynamics under a cost-free gene-for-gene model (GFG2, *d* = 0).** Without fitness costs on resistance or viral escape, resistance alleles sweep to fixation rather than cycling.

**S5 Fig. Correlations between transmission rate and host fitness components.** Scatter plots of transmission rate against relative hatch rate and relative eggs laid, shown separately for each sex.

**S6 Fig. Parameter space summary statistics for r = 0.27.** Heatmaps of coevolutionary outcomes and key statistics across infection cost, resistance cost, and transmission rate combinations.

**S7 Fig. Parameter space summary statistics for r = 0.70.** As S6 Fig, for a stronger resistance effect.

**S8 Data. Experimental data.** Excel spreadsheet containing all raw experimental data. Sheets contain: (1) summary statistics per host–virus genotype combination; (2) raw qRT-PCR data for viral load quantification; (3) raw transmission experiment data; (4) raw reproductive fitness assay data. (XLSX)

**S9 Data. Parameter sweep simulation output.** Summary statistics for all simulation runs in the two-virus gene-for-gene parameter sweep. Each row corresponds to one simulation run. Statistics prefixed “prevalent_” are calculated only over generations where both viruses independently exceed a 0.01% prevalence threshold. (XLSX)

## Acknowledgments

We thank Darren Obbard (University of Edinburgh) for assistance with wild fly collection, which yielded the UK Sigma virus isolates used in this study.

